# Mesoscopic Liquid Clusters Represent a Distinct Condensate of Mutant p53

**DOI:** 10.1101/2020.02.04.931980

**Authors:** David S. Yang, Arash Saeedi, Aram Davtyan, Mohsen Fathi, Mohammad S. Safari, Alena Klindziuk, Michelle C. Barton, Navin Varadarajan, Anatoly B. Kolomeisky, Peter G. Vekilov

**Affiliations:** Department of Chemical and Biomolecular Engineering, University of Houston, 4726 Calhoun Road, Houston, TX 77204-4004, USA; Department of Chemistry, Rice University, P.O. Box 1892, MS 60, Houston, TX 77251-1892, USA; Department of Molecular Biology, Princeton University, Princeton, NJ 08544-1014, USA; Department of Epigenetics and Molecular Carcinogenesis, The University of Texas MD Anderson Cancer Center, Houston, TX, 77030, USA; Department of Chemistry, University of Houston, 3585 Cullen Blvd., Houston, TX 77204-5003, USA

## Abstract

The oncogenic properties of mutant p53 have been ascribed to destabilization of the p53 conformation, followed by aggregation into insoluble fibrils. Here we combine immunofluorescent 3D confocal microscopy of breast cancer cells expressing the p53 mutant Arg248Gln (R248Q) with light scattering from solutions of the purified protein and molecular simulations to probe the mechanisms that govern phase behaviors of the mutant across multiple length scales, from cellular to molecular. We establish that p53 R248Q forms mesoscopic protein-rich clusters, an anomalous liquid phase with several unique properties. We demonstrate that the clusters host and facilitate the nucleation of amyloid fibrils. The distinct characteristics of the clusters of R248Q and wild-type p53 and theoretical models indicate that p53 condensation into clusters is driven by the structural destabilization of the core domain and not by interactions of its extensive disordered region. Two-step nucleation of mutant p53 amyloids suggests means to control fibrillization and the associated pathologies through modifying the cluster behaviors. In a broader context, our findings exemplify interactions between distinct protein phases that activate complex physicochemical mechanisms operating in biological systems.

## Introduction

The wild type p53 protein is a potent cancer suppressor, which is inactivated in almost every tumor, either through mutations in the TP53 gene (in 50% or more of human cancers) or deregulation of its associated pathways.^1–6^ By contrast, p53 mutants emerge as potent cancer promoters because they exert a dominant-negative (DN) effect on the wild-type variant and also display oncogenic gain-of-function (GOF) properties by inhibiting other cancer suppressors.^1^ Several mechanisms of cancer promotion by mutant p53 have been discussed.^6–12^ It was recently suggested that the structural destabilization of the p53 mutants and the associated aggregation into insoluble, degradation resistant structures, designated fibrils, may play a decisive role in their oncogenicity.^2,13–15^ Concurrently, fibril suppression, for instance, by stabilization of the mutant p53 conformation, has been identified as a general way to fight cancer.^2,16^

A parallel development identified a new form of biological organization, liquid-liquid phase separation, which constitutes several common membraneless organelles such as nucleoli, Cajal bodies, and P granules.^17–21^ Many proteins forming dense liquid condensates in live cells incorporate significant disordered regions and condensation at the low concentrations of these proteins in the nucleoplasm and cytoplasm has been attributed to stronger intermolecular attraction due to the unstructured segments.^19,20^ Furthermore, it is well-appreciated that protein dense liquids can serve as precursors and facilitators of higher order organization such as ordering into crystals ^22–25^ and the formation of polymers of sickle hemoglobin.^26,27^ Theoretical analyses and recent experiments suggest that a similar precursor mechanism would apply to the nucleation of amyloid fibrils.^28–33^

Here we explore whether dense liquid phases may form in mutant p53 solutions and facilitate the nucleation of fibrils. We monitor the phase behaviors of p53 R248Q in a cell line expressing this mutant and in solutions of the purified protein. As p53 binds to DNA, the positive arginine (R) 248 residue embeds in the minor grove of the double helix to support the contact (Figure 1a).^2,34^ Mutations at this site are the most frequent oncogenic p53 mutations.^2,34^ Besides weakening the binding to DNA (Figure 1a), replacing arginine with glutamine (Q) destabilizes the structure of the DNA binding domain and strongly enhances the aggregation propensity of p53.^2,34^ The combination of weaker binding and structural destabilization classifies R248Q among the strongest predictors of patient death in ovarian cancer.^35^

**Figure 1.**
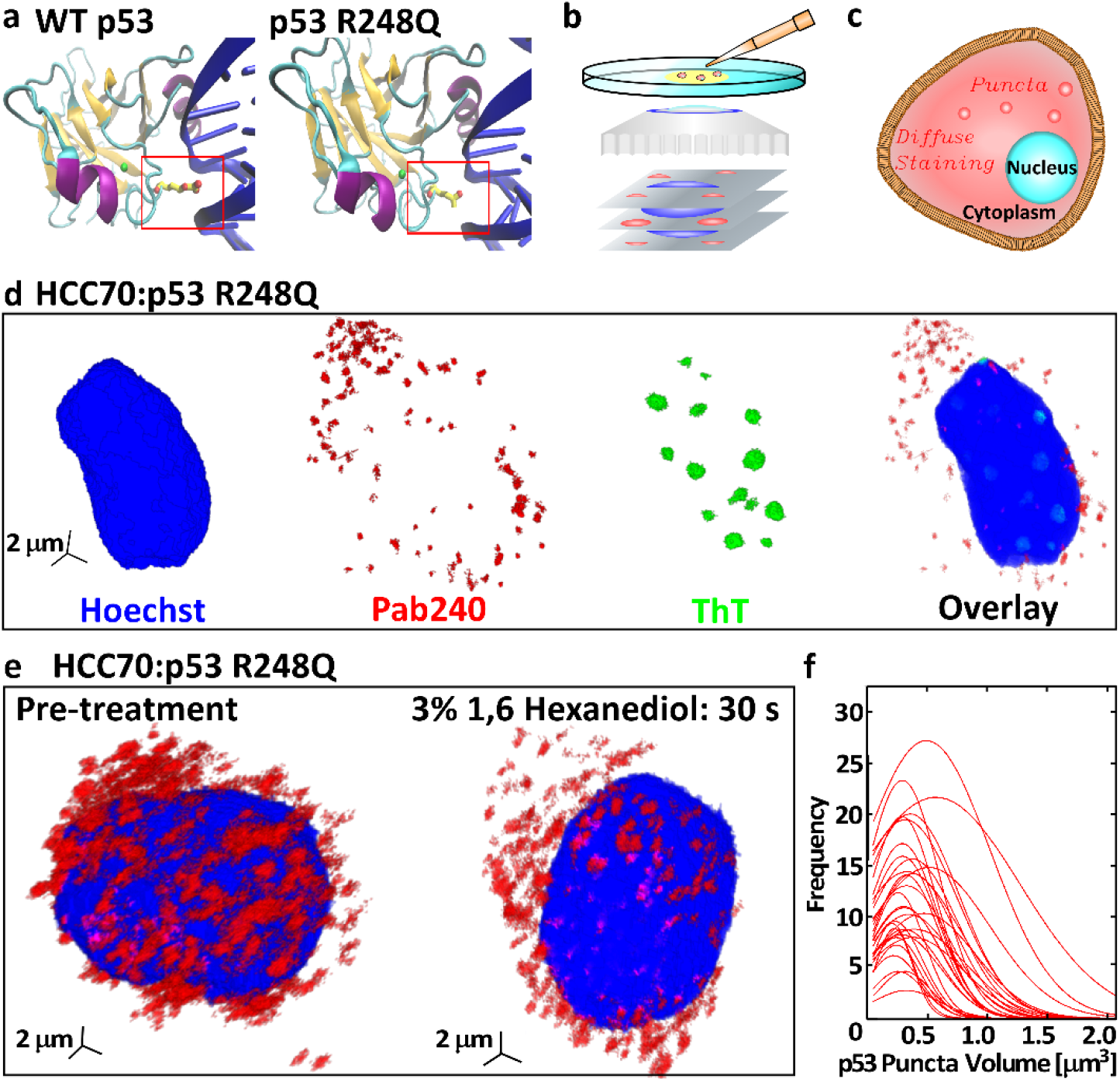
The R248Q mutation and aggregation on p53 R248Q in breast cancer cells. **a.** The structure of the DNA-binding domain (DBD, 94-292) of wild-type p53 and the R248Q (Arg 248 Gln) mutant. Nitrogen atoms are colored in red, zinc in green, alpha-helices in purple, beta sheets in orange, and DNA in blue. PDB ID: 1TUP ^40^ for wild type p53; prediction of p53 R248Q DNA interaction using VMD.^41^ **b.** Schematic of confocal immunofluorescent microcopy, in which antibodies that specifically target cell components of interest are tagged with fluorescent dyes. The spatial distribution of the fluorescent signal collected by the microscope maps the 3D distribution of the target molecule. **c.** Illustration of diffuse and punctate staining in the cytoplasm. **d.** Combined staining with Pab240, which binds to unfolded or aggregated p53, and ThT, which detects amyloid structures. The cell nucleus is stained with a Hoechst dye. **e.** Staining of HCC70 cells with Pab240 before and after treatment with 1,6-hexanediol, known to disperse dense liquid droplets of disordered proteins. **f.** Distributions of the volumes of the puncta found in HCC70 cells treated with Pab240. Each trace represents the volume distribution of puncta from a single cell.

We examine the existence of two distinct liquid phases: macroscopic liquid, similar to the phase found with other partially disordered proteins,^17–21^ and mesoscopic protein-rich clusters observed with numerous globular proteins ^23–25,36,37^ and recently found for wild type p53.^38^ We establish that p53 R248Q forms mesoscopic clusters, which are distinct form the macroscopic dense liquid, and that they host the nucleation of amyloid fibrils. The proposed mechanism of fibrillization facilitated by liquid condensates drastically deviates from the accepted sequential association of single solute molecules. We define the molecular mechanism of the enhanced aggregation of the p53 mutants. Comparison of the thermodynamic and kinetic characteristics of condensation of R248Q and wild-type p53 and theoretical models indicate that p53 condensation and aggregation are driven by the destabilization of the core domain and not by interactions of its extensive disordered region.^34^ This finding is consistent with the localization of most cancer-associated mutations in the structured DNA binding domain.^15^

## Results

### Cytoplasmic aggregation of P53 R248Q in cancer cells

We explore the phase behaviors of p53 in two breast cancer cell lines, HCC70, which expresses P53 R248Q and MCF7, expressing wild type p53,. We detect and quantify aggregated and unaggregated p53 by immunofluorescent 3D confocal microscopy (Figure 1b), which exploits the sensitivity of antibodies to their antigen to attach fluorescent dyes to specific targets within a cell and map the 3D distribution of the target molecule.^39^ Uniformly distributed targets present diffuse staining, whereas aggregates of target molecules appear as puncta (Figure 1c). To identify the nucleus, each cell is treated with a Hoechst dye, which binds to DNA and emits a characteristic blue signal.

To detect and characterize p53 aggregates, we perform three-dimensional multi-color immunofluorescent staining. We combine staining with an antibody specific for misfolded or aggregated p53, Pab240, and Thioflavin T (ThT), a common probe for amyloid structures. HCC70 cells exhibit exclusively cytosolic, punctate Pab240 staining and lack of detectable p53 staining in the nucleus (Figures 1c and S2). ThT staining is pronounced in the nucleus, where it indicates the presence of non-53 amyloid formations; no ThT staining is detectable in the cytoplasm (Figure 1c). Taken together, the two staining patterns reveals the existence of aggregated p53 R248Q within the cytoplasm of HCC70 cells and suggest that the aggregates are not p53 fibrils. Treating the HCC70 cells with the antibody DO1, directed against the N-terminal transactivation domain of p53, reveals diffuse staining in the cell nucleus (Figure S1a), indicating an elevated, compared to the cytoplasm level, concentration of unaggregated p53 in the nucleus of HCC70 cells. By contrast, MCF7 cells (expressing wild type p53) treated with the PAb240 antibody displayed only strong perinuclear staining (Figure S3). The protein accumulated in vicinity of the nucleus appears stable and unaggregated since it also stains with the DO1 antibody (Figure S1b).

The lack of ThT staining of the p53 R248Q aggregates in HCC70 cells suggests that the aggregates are not amyloid fibrils. To test whether the aggregates are droplets of macroscopic dense liquid, we determined the sensitivity of the cellular puncta to treatment with 1,6-hexanediol, an organic molecule known to destabilize liquid condensates.^18,42^ HCC70 cells treated with 1,6-hexanediol showed no reduction in the number of cytoplasmic p53 puncta (Figure 1f), indicating the p53 R248Q aggregates are not dense liquid droplets. We measured the volume of the individual puncta of p53 R248Q in the HCC70 cells. The size distributions of 32 cells are similar and relatively narrow with average volumes ranging from 0.1 to 0.6 μm^3^ (Figure 1e). The reproducible narrow and weighted to small sizes volume distributions of the aggregates are not consistent with behaviors expected for disordered agglomerates, whose stochastic formation at low driving forces may result in broader distributions and greater variability between cells.

Collectively, imaging with Pab240 and DO1 antibodies and ThT, and the response to 1,6-hexanediol demonstrate that the mutant p53 R248Q forms aggregates of narrow size distributions within the cytoplasm of breast cancer cells whereas wild type p53 does not aggregate within cancer cells. The results with breast cancer cells establish that the p53 R248Q protein aggregates within the cytoplasm are not fibrils or droplets of stable dense liquid. For further insight into the mechanisms and properties of the observed aggregates, we turn to experiments under defined conditions *in vitro*.

### Mesoscopic protein-rich clusters in solutions of p53 R248Q

We monitored solutions of p53 R248Q with oblique illumination microcopy (OIM, Figure 2a).^43,44^ This method records speckles of light scattered by individual solution inhomogeneities and is particularly suited to detection of mesoscopic aggregates since, according to the Rayleigh law, the scattered light intensity scales with the sixth power of the scatterers’ size. The solutions were filtered through low-protein binding 220 nm filters to remove extrinsic inhomogeneities and loaded on the microscope within 10 – 20 min of preparation. The OIM micrographs reveal speckles of light that correspond to protein aggregates. Notably, the aggregates are not amyloid structures since tests using the fluorescent dye anilinonaphthalenesulfonate (ANS), discussed below, reveal that at concentrations similar to the one employed here, 2 μM, amyloid fibrillization is delayed by hours.

**Figure 2.**
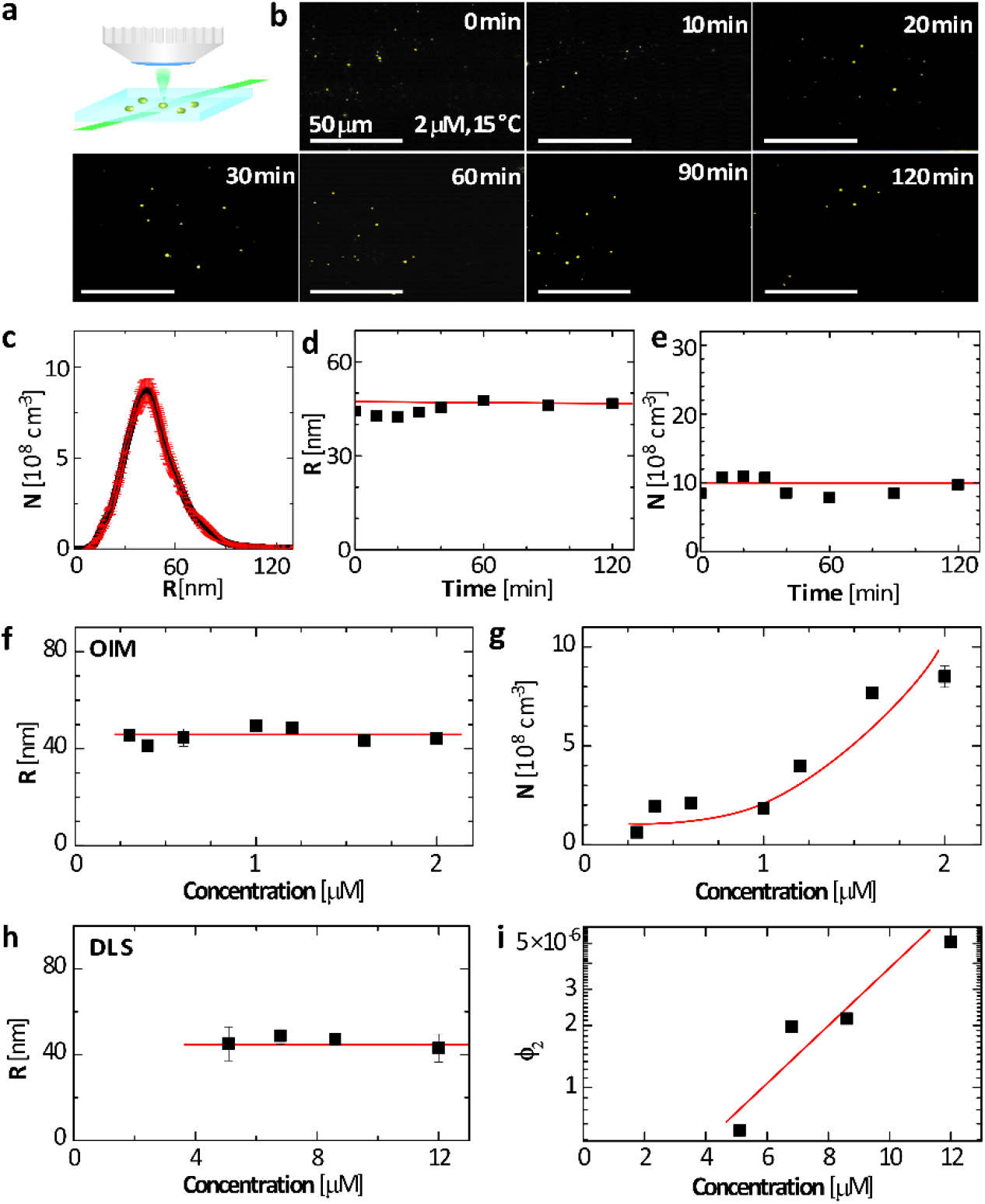
Mesoscopic protein-rich clusters of p53 R248Q. **a.** Schematic of oblique illumination microscopy (OIM). A thin solution is illuminated by green laser at an oblique angle. Scattered light is collected by a microscope lens **b.** Representative OIM micrographs tracing the evolution of a 2 μM P53 R248Q solution at 15 °C. The observed volume is 5 × 80 × 120 μm^3^ depth × height × width. The clusters appear as gold speckles. **c.** Number density distribution of the radii R of clusters determined by OIM. The averages of five measurements are displayed. The error bars represent the respective standard deviations. **d. e.** The evolution of the average radius *R* and number concentration *N* of clusters determined at 15°C by OIM from images as in **b.** The averages of five measurements are displayed. The respective standard deviations are smaller than the symbol size. Horizontal lines denote the mean values of *R* and *N*. **f, g.** The concentration dependence of R and N determined at 15°C by OIM. The averages of five measurements are displayed. The error bars represent the respective standard deviations and are smaller than the symbol size for most determinations. Horizontal line in **f** denotes the mean value of *R*; curve in **g** is a guide to the eye. **h, i.** The concentration dependence of R and the cluster volume fraction ϕ_2_ determined at 15°C by dynamic light scattering. The averages of five measurements are displayed. The error bars represent the respective standard deviations and are smaller than the symbol size for some determinations. Horizontal line in **h** denotes the mean value of *R*; line in **g** is a guide to the eye.

To distinguish the aggregates observed with p53 R248Q from macroscopic dense protein liquids and amorphous agglomerates, we monitor the evolution of their radii *R* and number concentrations *N* and the correlations of *R* and *N* with the protein concentration. We determine the aggregates’ radii from their Brownian trajectories, extracted from sequences of OIM images.^43,44^ The aggregates exhibit a relatively narrow size distribution (Figure 2c) with an average of R = 45 ± 5 nm at 2 μM and 15°C. Both *R* and *N* are steady for at least two hours (Figure 2d,e), behaviors that stand in contrast to expectations for liquid-liquid separation, a first-order phase transition,^45,46^ for which nucleation of new liquid droplets and their growth persist and *R* and *N* increase.^47,48^ The reversibility of the observed aggregates is revealed by the concentration dependence of their number concentration (Figure 2g) and the fraction of the solution volume that they occupy ϕ_2_ (Figure 2i). The concentration *N* declines from 8.7×10^8^ cm^-3^ to 0.4×10^8^ cm^-3^, about 20-fold, in response to a seven-fold reduction of concentration, from 2.0 to 0.3 μM. Similarly, ϕ_2_, determined independently by dynamic light scattering (DLS), shrinks from 5×10^-6^ to 0.5×10^-6^, a ten-fold decrease driven by 2.5-fold lower concentration. The two techniques complement their respective concentration ranges and demonstrate that *R* is consistently about 45 nm at *C_0_* between 0.2 and 12 μM. The OIM measurement of *N* is consistent with the DLS determination of ϕ_2_: the product *4nR^3^N/3* = 0.3×10^-6^ at 2 μM is close to the ϕ_2_ value extrapolated for that concentration. The exaggerated response of *N* and ϕ_2_ to reduced concentration indicates that the aggregates are not irreversible disordered agglomerates, whose concentration is diluted in parallel with that of the protein, but rather condensates existing in dynamic equilibrium with the host solution.

Three behaviors of the p53 R248Q aggregates cohere with previous observations of mesoscopic protein-rich clusters of both globular proteins and wild type p53.^38,43,49,50^ The average cluster radius *R* is, first, steady and, second, independent of the protein concentration *C_o_*. The third characteristic cluster behavior exhibited by the p53 R248 aggregates is the near-exponential increase of *N* and ϕ_2_ with *C_o_* (Figure 2 h,j). We conclude that the aggregates are mesoscopic protein-rich clusters. According to recent models the mesoscopic liquid clusters of multi-chain proteins, such as tetrameric p53, exist due to the accumulation of misassembled oligomers.^38,51,52^ The cluster size is determined by the balance between the lifetime of the misassembled oligomers and their rate of outward diffusion from the cluster core and is, hence, independent of the protein concentration and steady in time.^51,53^ By contrast, the amount of protein captured in the clusters, and the related number of clusters and cluster population volume, increases exponentially with the protein concentration as a consequence of the thermodynamic equilibrium between the clusters and the bulk solution.^37,51,53^ The mesoscopic clusters of both R248Q and wild type p53^53^ appear to remarkably well comply with the predictions of this model.

### The enlarged aggregation capacity of p53 R248Q

To understand how the R248Q mutation impresses the behaviors of the mesoscopic clusters we explored the response of the cluster characteristics to temperature. Solution samples were incubated for 20 min at five temperatures between 15 and 42°C. OIM observations revealed that the clusters of P53 R248Q are sensitive to temperature variation (Figure 3a – c). Increasing the temperature from 15°C, discussed above, to 18, 25, 37 and 42°C increases cluster formation (Figure 3a), in concert with previous observation with wild type p53.^38^ The mid-denaturation temperature of p53 R248Q is 37°C,^54^ however at the tested concentration, 2 μM, fibrillization is significantly delayed, as discussed below. The greater cluster number at elevated temperatures suggests that unfolding of the p53 R248Q core domain may be an essential trigger for cluster formation.

**Figure 3.**
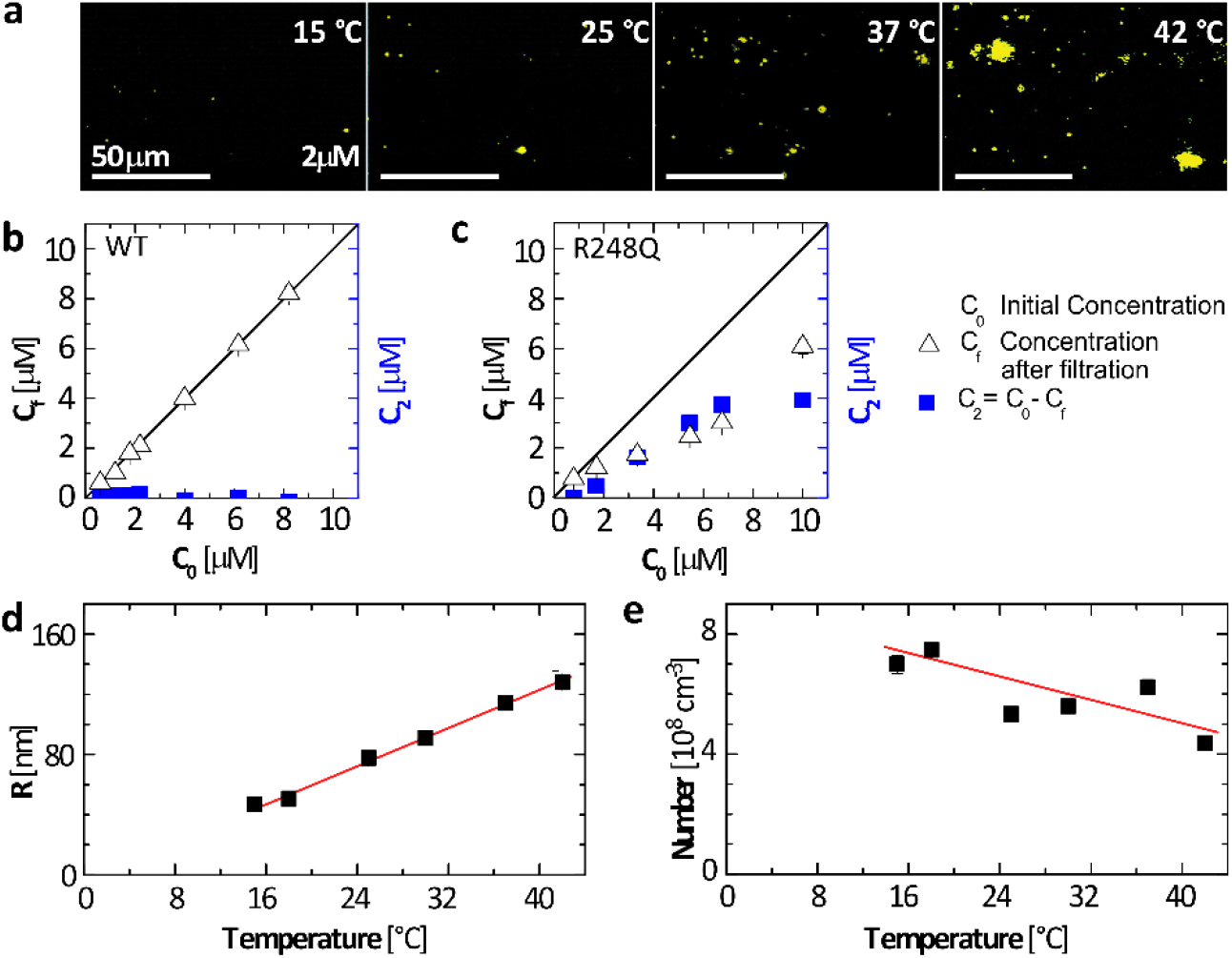
p53 R248Q clusters at temperatures between 15 and 42°C. **a.** Representative OIM micrographs collected from a p53 R248Q solution with concentration C_0_ = 2. μM after incubation for 20 minutes at each temperature. The observed volume is 5 × 80 × 120 μm^3^ depth × height × width. Aggregates appear as cyan speckles. **b, c.** Concentrations *C_f_* and *C*_2_, defined in the plot, of wild type p53, in **b**, and P53 R248Q, in **c**, after incubation for 20 min at 15°C as a function of the initial solution concentration *C_0_*. ***d, e*.** The average radius R and the total number N, respectively, of the aggregates, determined by OIM from images as in **a**. The average of five determinations in distinct solution volumes is shown. Error bars indicate standard deviations and may be smaller than the symbol size. Lines are just guides to the eye.

p53 R248Q manifests an enlarged capacity to form clusters, exposed by cluster formation at 15°C and *C_o_* = 2 μM (Figure 3a), in contrast to wild type p53, which exhibits no clusters at these temperature and concertation.^38^ This exaggerated cluster formation is reaffirmed by tests at *C_o_* as high as 8 μM. The concentration of the solution in equilibrium with the clusters, *C_f_*, measured after removing the clusters by filtration, is equal to the initial *C_o_* with wild type p53 (Figure 3b), indicating that no protein is captured in clusters or any other aggregates; this observation conforms to the lack of clusters in the OIM tests.^38^ By contrast, filtration to remove the clusters in a p53 R248Q solution lowers *C_f_* from *C_o_* by about half (Figure 3c), indicating that that the clusters hold about 50% of the dissolved mutant. The correlation between *C_f_*, which represents the concentration of equilibrium between the clusters and the solution, and *C_o_* is an additional distinguishing characteristics of the mesoscopic clusters. This correlation is striking contrast with examples of dense protein liquids and amyloid fibrils; these two aggregate classes equilibrate with solutions of constant concentration, referred to as solubility.^19,20,27,55–57^ The correlation between *C_f_* and *C_2_* is likely represented by a thermodynamic model relying on the complex composition of the phase captured in the clusters, originally developed for the mesoscopic clusters of wild type p53.^38^

The cluster radius *R* increases with temperature and reaches ca. 130 nm at 37°C. This size approaches the value ca. 300 nm, corresponding to the average volume *V_c_* ≅ 0.1 μm^3^ of the p53 R248Q aggregates observed in HCC70 cells, according to the relation *V_c_* = 4*πR*^3^/3. In view of the enlarged clustering capacity of the p53 R248Q, it is surprising that the average *R* increases linearly with temperature and the number of clusters *N* weakly declines (Figure 3d,e). These trends are in contrast to the *R*(*T*) and *N*(*T*) correlations for wild type p53, both of which are increasing and exponential.^38^ The abridged periods between solution preparation and the *R* and *N* measurements suggest that this unusual response to increasing temperature is not due to concurrent growth of alternative aggregates (fibrils, disordered agglomerates, etc.) that capture the majority of the protein and constrain cluster formation.

To understand the response of *R* and *N* of the p53 R248Q clusters population to temperature, we examine the oligomer state of the unaggregated protein in equilibrium with the clusters. The correlation functions *g*_2_(τ) of the light scattered from p53 R248Q solutions at three concentrations reveal the presence of two distinct scatterers with characteristics diffusion times τ_1_ and τ_2_ (Figure 4a). The longer diffusion times τ_2_ correspond to the slower diffusion of larger scatterers, the clusters. The shorter τ_1_s represent the faster diffusion of the unaggregated protein. Notably, the *g_2_* shoulder with τ_2_ is faint, indicating that the majority of the light is scattered by unaggregated protein. The cluster radii *R* (Figure 2i) computed from the τ_2_s are agree with the OIM measurements (Figure 2g). The average radii *R_u_* of the unaggregated protein, evaluated from the τ_1_s, range from 8 to 12 nm (Figure 4b), substantially greater than the ca. 4 nm radius expected for a p53 tetramer of molecular weight 175 kg mol^-1^.^58^ Consonantly with the exaggerated radius, the considerable polydispersity μτ^2^ ≈ 0.35 – 0.25 (Figure 4c) indicates the dominance of high-order oligomers over monomeric, dimeric, and tetrameric p53 units.^14,59^

**Figure 4.**
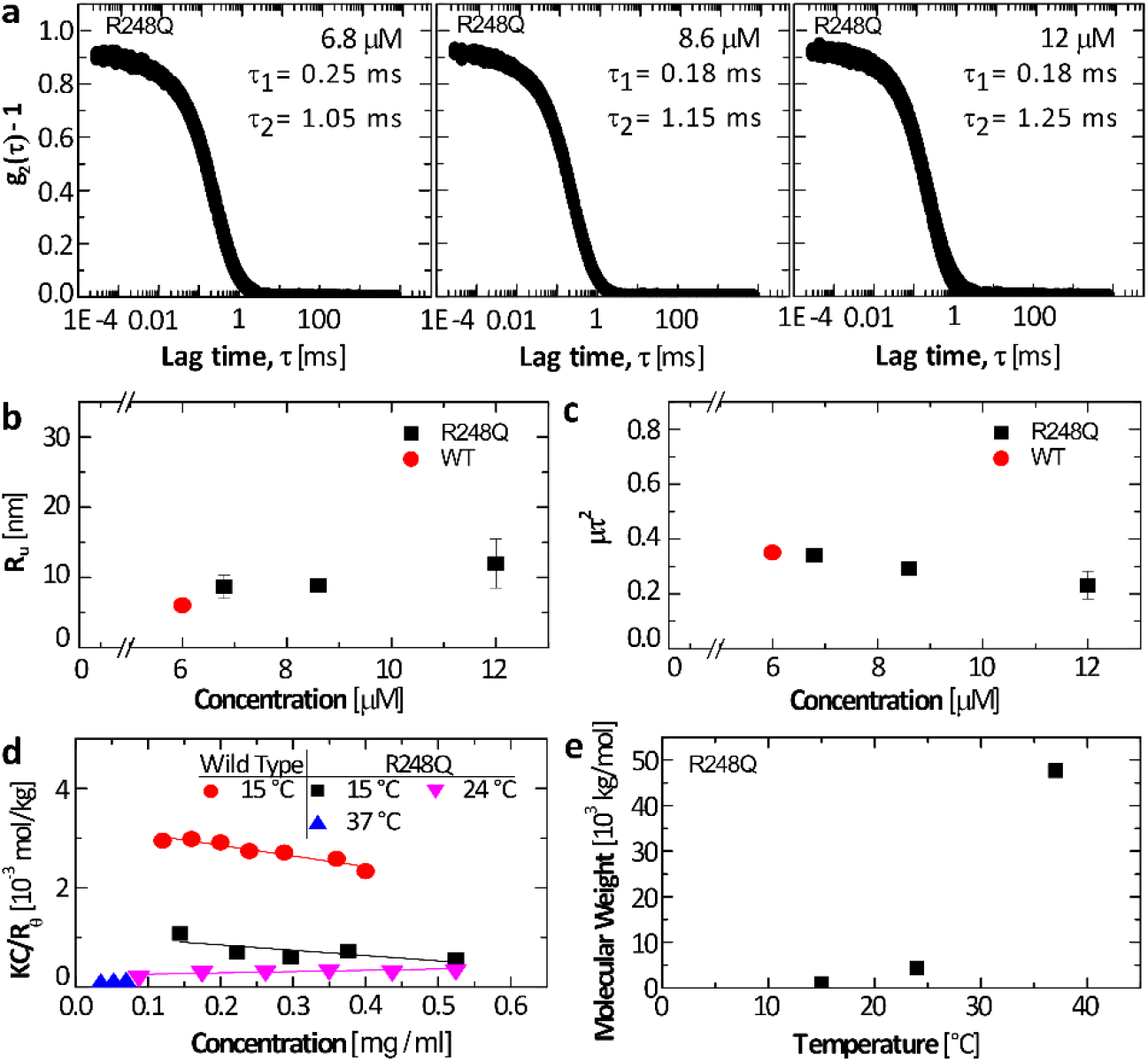
The structure of unaggregated p53 R248Q. **a.** Normalized intensity correlation functions *g_2_*(τ) of the light scattered by filtered p53 R248Q solutions at shown concentrations at 15°C. The characteristic diffusion times of unaggregated protein τ_1_ and of the τ_2_ clusters τ_2_ are shown. **b, c.** Average radius *R_m_* and dispersity μτ^2^ of unaggregated p53 R248Q determined from the *g*_2_s in **a.** for p53 R248Q. Data for wild type p53^38^ are shown for comparison. **d.** The osmotic compressibility of the solution *KC*/*R_θ_*(*K*, instrument constant; *R_θ_*, Rayleigh ratio of the intensity of light scattered at 90° to the incident light; C, concentration of p53) as a function of concentration for wild type p53 at 15 °C and R248Q at 15 °C, 24 °C, and 37 °C. **e.** The average molecular weight of unaggregated p53 R248Q evaluated form the *KC*/*R_θ_*(*C*) data in d at three temperatures.

To further explore how temperature manipulates the state of the unaggregated protein, we measured average protein molecular weight *M_w_* using static light scattering at 15, 24, and 37°C (Figure 4d,e). The intercepts of the linear correlations *KC*/*R_θ_*(*C*) (*K*, instrument constant; *R_θ_*, Rayleigh ratio of the intensity of light scattered at 90° to that of the incident light; *C*, concentration of p53) (Figure 4d)^60^ equal the reciprocal *M_w_* of the unaggregated protein. At 15°C, *M_w_* ≅ 943 kg mol^-1^ for P53 R248Q, which is about three-fold greater than that of wild type p53 and five-fold higher than that of the p53 tetramer with *M_w_* = 175 kg mol^-1^. The exaggerated molecular weight coheres with the inflated monomer size *R_u_* and high polydispersity μτ^2^ determined by dynamic light scattering (Figure 4a – c). As temperature increases, *Mw* grows to 4500 kg mol^-1^ at 24°C and 48,000 kg mol^-1^ at 37°C indicating excessive oligomerization of p53 R248Q. Collectively, the static and dynamic light scattering characterizations of the unaggregated protein reveal the presence of high order oligomers that coexist with the mesoscopic clusters. At elevated temperatures, both the average size of the oligomers and amount of protein sequestered in them increase. As a result, the amount of protein available to the clusters abates, driving down the motivation for cluster formation and the number of clusters.

### The mesoscopic protein-rich clusters host the nucleation of p53 R248Q fibrils

Formation of amyloid fibrils of mutant and wild type p53 is a distinguishing behavior of this protein.^14,61,62^ To examine whether the mesoscopic protein-rich clusters appertain to the mechanisms of nucleation and growth of the P53 R248Q fibrils we employed as a probe the response of the fibrillization kinetics to Ficoll. We monitored the growth of the amyloid population with the dye 1-anilino-8-naphthalenesulfonate (ANS), which binds to amyloid fibrils and emits fluorescence at 500 nm.^63^ Notably, ANS also binds to exposed hydrophobic regions abundant in partially unfolded proteins;^63^ the pronounced fluorescence intensity in both wild type and mutant p53 solutions immediately after ANS introduction (Figure 4a,b) may be due to the binding of the dye to the disordered segments in the transactivation and proline-rich domains.^2^ The stronger initial fluorescence of the mutant solution attests to the abundance of hydrophobic residues exposed owing to the lower stability of its core domain.^2,64^

In the absence of Ficoll solutions of wild type p53 emitted steady fluorescence intensity for ca. 40 min at the highest tested concentration, 6.5 μM, and for up to nine hours at the two lower concentrations (Figure 5a). After this lag time, the intensity ascended. The observed fibrillation delay is likely due to the known slow nucleation of amyloid structures.^13,65^ The R248 p53 at 6.5 μM fibrilizes after a shorter lag time, ca. 20 min (Figure 5b), indicating fast nucleation of the mutant fibrils; at the lower tested concentrations, the nucleation of R248 p53 fibrils is delayed by at least 5 hours (Figure 5b).

**Figure 5.**
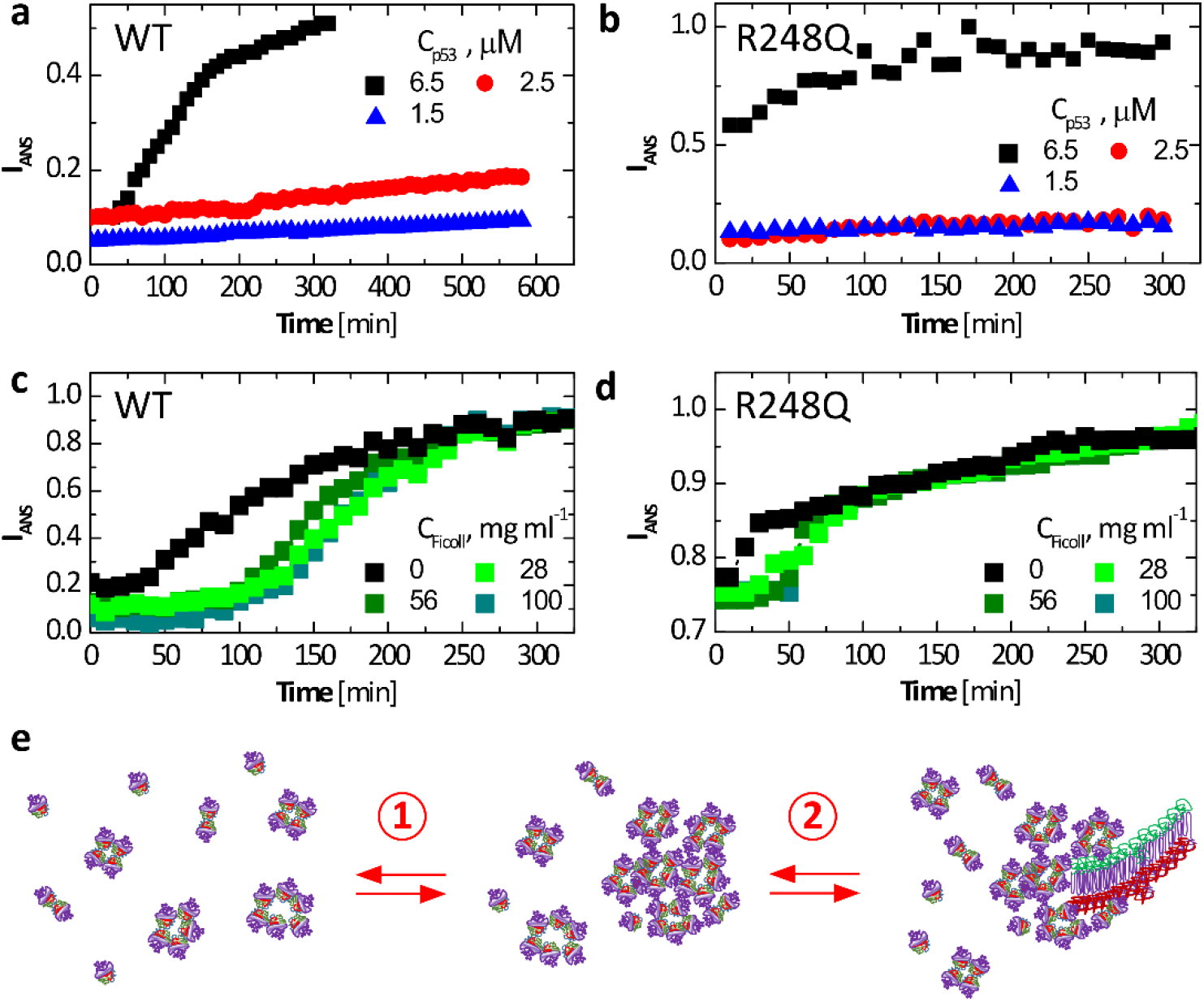
Fibrillization of wild type and p53 R248Q. **a – d.** Evolution of the intensity of fluorescence at 500 nm of 1-anilino-8-naphthalenesulfonate (ANS) in the presence of p53 at 37°C. ANS concentrations was 200 μM in all tests. **a, b.** At the listed concentrations of wild type, in **a**, and R248Q, in **b**, in the absence of Ficoll. **c, d.** At 6.5 μM of wild type, in **c**, and R248Q, in **d**, and in the presence of varying concentrations of Ficoll. **e.** Schematic of two-step nucleation of fibrils. Step 1. Mesoscopic p53-rich clusters form from misassembled p53 oligomers and native tetramers. Step 2. Fibrils nucleate within the mesoscopic clusters. Fibril growth proceeds classically, via sequential association of p53 monomers from the solution.

Time-dependent ANS fluorescence reveals that added Ficoll invokes significantly longer lag times with both wild type and p53 R248Q (Figure 5c,d). Ficoll-enforced nucleation delay is counterintuitive since the excluded volume effects of Ficoll and the associated surge of the protein chemical potential^66^ would hasten fibril nucleation. The faster fibril growth in the presence of Ficoll, manifest as steeper gain of ANS fluorescence after the lag time for both wild type and mutant p53 (Figure 5 c,d), concurs with crowding-enforced chemical potential boost. On the other hand, suppressed nucleation in the presence of Ficoll coheres with nuclei growth hosted within the clusters. Ficoll sequesters in the clusters,^38^ where it may obstruct the migration of the p53 molecules to a fibril nucleus. The accelerated fibril growth in the presence of Ficoll suggests that after nucleation, the fibrils emerge from the clusters and grow in the p53 solution. The emerging non-classical two-step mechanism of fibril nucleation assisted by preformed mesoscopic clusters, and followed by classical growth by association of solute monomers, is illustrated in Figure 5e.

### Why is R248Q mutant more prone to aggregate than the wild type?

To understand how a mutation located in the ordered DNA binding domain (DBD, comprised of residues 94 to 289) of p53 lowers the stability of the molecule and promotes aggregation we modeled the conformational changes in the p53 structure driven by the R248Q mutation. We use the Associative Memory, Water Mediated, Structure and Energy Model for Molecular Dynamics (AWSEM-MD).^67^ For a coarse estimate of the conformational modifications enforced by the mutation, we first evaluate the free energy profiles *F* of the wild type and P53 R248Q DBDs.^67^ We introduce a reaction coordinate *q* that measures the similarity of the DBD conformation to an experimentally known DBD structure (*q* = 0 for random coils; *q* = 1 for structures identical to the protein database entry). Notably, the experimental structure of reference used here is of the DBD bound to DNA,^40^ which might be significantly different from the unbound structure that we model. Indeed, we found that the free energy minima for both wild type and p53 R248Q locate at *q* values below 0.7; this value indicates a high degree of similarity between structures; *q* < 1 manifests the divergence between the DNA-bound and unassociated structures. Importantly, the 1D free energy profile *F*(*q*) reveals that wild type and p53 R248Q explore distinct conformational spaces (Figure 6a). Whereas the free energy minimum for the wild type is at *q* > 0.5, it is closer to 0.45 for the R248Q mutant.

**Figure 6.**
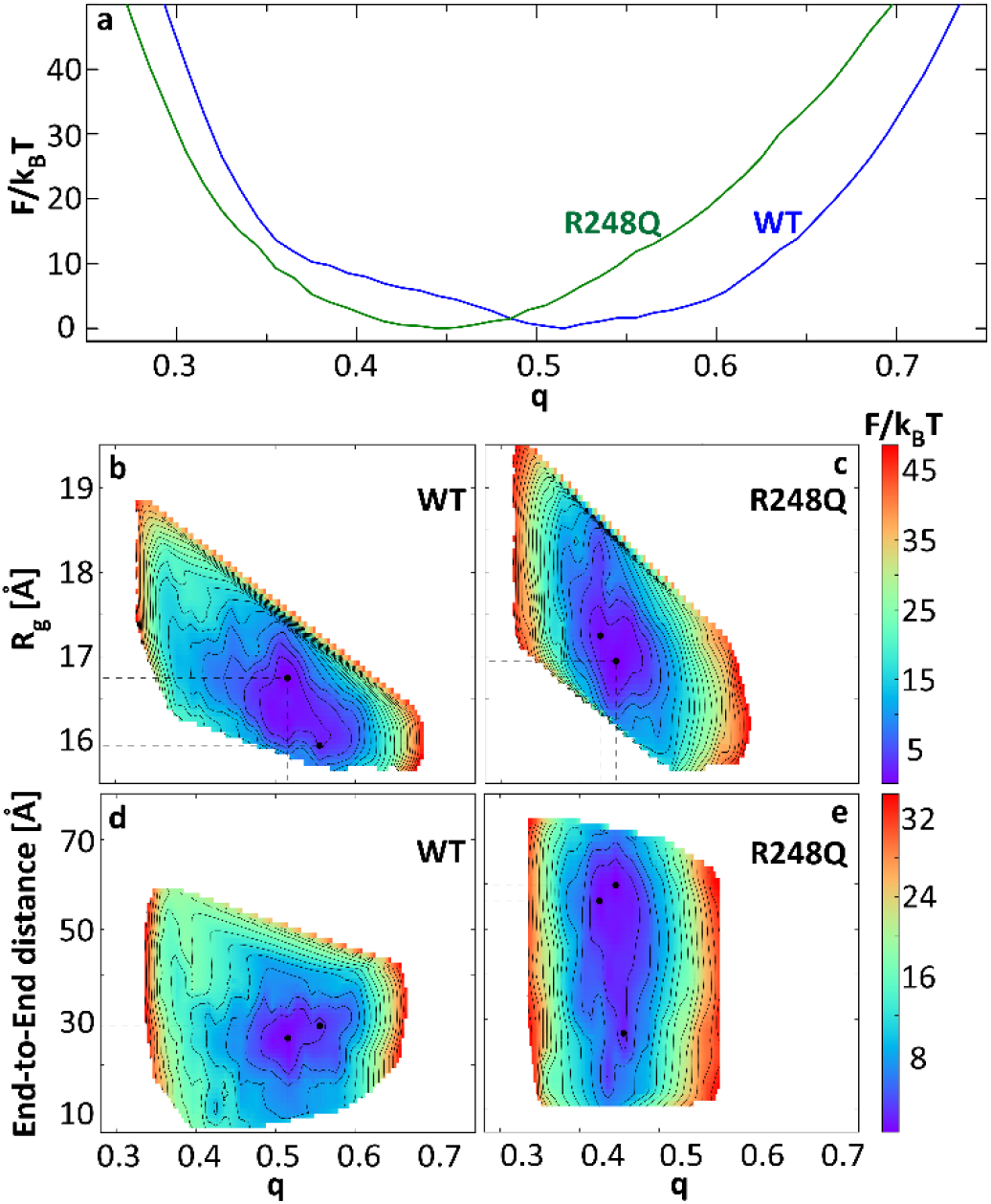
The free energy *F* of the conformations of wild type and p53 R248Q DNA binding domains. **a.** As a function of the reaction coordinate *q* that measures the similarity of the DBD conformation to a structure determined by x-ray crystallography.^40^. **b – e.** Two dimensional profiles of *F* as function of *q* and radius of gyration, *R_g_*, in **b** and **c**, and end-to-end distance, *D*, in **d** and **e**, of the DBD chain

For further insight, we explore the free energy as a function of two additional metrics of protein confirmation: the radius of gyration, *R_g_*, and end-to-end distance, *D*, of the DBD chain. The 2D free energy profiles *F*(*q*, *R_g_*) and *F*(*q*, *D*) reveal that there are at least two local minima with *F* below *k_B_T*(*k_B_*, Boltzmann constant; *T*, temperature) for both wild type and mutant p53 (Figure 6b – e). For wild type p53, *F*(*q*, *R_g_*) minima locate at at *q* = 0.515 and *R_g_* = 16.75 Å and at *q* = 0.555 and *R_g_* = 15.95 Å (Figure 6b). For the mutant, the minima are at *q* = 0.425 and *R_g_* = 17.25 Å and *q* = 0.445 and *R_g_* = 16.95 Å (Figure 6c). Thus, the mutant explores structures characterized with lower *q* and higher *R_g_*. Consistently, the *F*(*q*, *D*) free energy profiles of the wild type exhibit two minima, at the same *q* as *F*(*q*, *R_g_*) and end-to-end distances 26 Å and 28.6 Å, both 30 Å (Figure 6d). For the mutant, however, we identify three local minima can be identified at *q* values 0.425, 0.445, and 0.455. While the minimum at *q* = 0.455 has end-to-end distance comparable to that of the wild type, ca. 27 Å, the minima at *q* = 0.425 and *q* = 0.445 have larger *D* values of 56 Å and 60 Å, respectively. The *F*(*q*, *R_g_*) and *F*(*q*, *D*) profiles indicate that the mutant DBD can adopt extended conformations that have less similarity to the reference DNA-bound structure than the wild-type DBD.

We model the wild type and mutant p53 DBD conformations near the free energy minima in the *F*(*q*, *R_g_*) and *F*(*q*, *D*) profiles. Pairwise comparisons of representative structures reveal that the conformations associated with two local minima of the *F*(*q*, *R_g_*) and *F*(*q*, *D*) profiles of the wild type are similar (Figure 7a). The largest difference is associated with a region containing a small helix and a large loop between residues 168 and 193. Both the N-terminal tail and the C-terminal helix, roughly defined as first and last 13 residues of DBD, are tightly wrapped around the domain at all times. Importantly, the *N*-terminal tail is bound to the ILTIITL motif (residues 251 to 257), known to promote p53 aggregation.^14,65^ The mutant structures are, however, very different. The largest changes are associated with the *N*-terminal and *C*-terminal tails that are both found in various unbound conformations (Figure 7b – d). The unbinding of the *N*-terminal loop and *C*-terminal helix can lead to significant changes to the full-length p53 structure and dynamics. Importantly, the unbinding of the N-terminal tail leads to the exposure of the 252ILTIITL257 motif known to promote p53 aggregation.^14^ The exposure of the aggregation prone sequence in the p53 R248Q may be a part of the molecular mechanism of enhanced cluster formation, oligomerization, and aggregation of this mutant.

**Figure 7.**
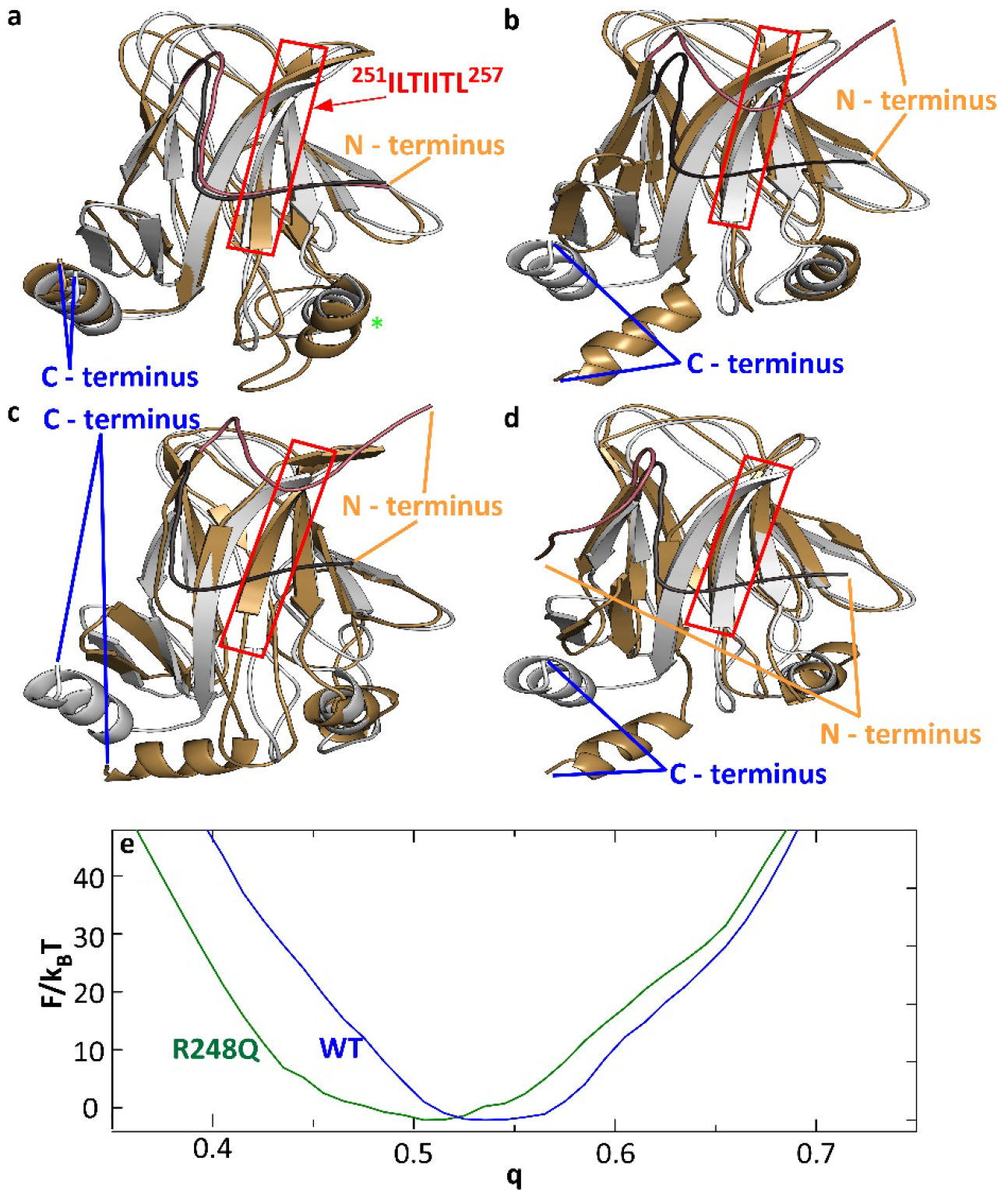
Conformational changes induced by the R248Q mutation. **a.** Comparison of wild type DBD structures corresponding to *F*(*q*) minima at q = 0.555 (silver) and q = 0.515 (gold). Red box highlights the aggregation-prone sequence, protected by the N-terminus tail in wild type p53 and exposed in p53 R248Q. Green star indicates residues 168 to 193, the location of the strongest deviation of between the two modeled wild type conformations. **b.** Comparisons of wild type DBD structure corresponding to the *F*(*q*) minimum at q = 0.555 (silver) to the DBD structures of p53 R248Q (gold) at *F*(*q*) minima at q = 0.425 in **b**; at q = 0.445 in **c**; and at q = 0.455 in **d**. In **a – d**, the N-terminus tail of the reference structure is highlighted in charcoal, and that of the second structure, in copper. **e.** Free energy profiles *F*(*q*) for the cores of the DBDs (residues 107 to 276) of wild type and p53 R248Q.

To understand how a mutation near the center of the DBD drives strong displacements of both DBD chain termini, we evaluated the *F*(*q*) profiles for the DBD core, comprised of residues 107 to 276 and omitting 13 residues from both N- and C-ends of the DBD, for both the wild type and the mutant. The *F*(*q*) minimum shifts from ca. 0.54 to ca. 0.51 (Figure 7e). The modified *F*(*q*) profile indicates that the mutation induces conformation changes in the DBD core, albeit more subtle than for the entire DBD (Figure 6a). The conformational changes in the DBD core illuminate how the structure perturbation introduced by the mutation propagates allosterically to the C- and N-termini and how the mutation drives the interactions of the α-helix at the C-terminus and the tail at the N-terminus with the core.

## Discussion

The mesoscopic protein-rich clusters of P53 R248Q, observed in cell culture and in solutions of purified protein represent a unique protein condensed phase. The defining features of the clusters are the decoupled cluster size and total cluster population volume, independence of the cluster size on the solution thermodynamic parameters, including the p53 concentration, and the variable solution concentration at equilibrium. The clusters share specific characteristics with the dense liquids observed with other proteins:^19–21^ their formation is reversible, they are in equilibrium with the solution, and they capture up to 80% of the available protein. The dramatic differences of their spatial and thermodynamic characteristics from those of protein dense liquid phases, however, certify the distinction of the clusters from dense liquid droplets.

The formation of p53 fibrils starts with nucleation, whereby local fluctuations of the p53 concentration beget regions of concentrated and ordered p53 molecules that serve as nuclei for the growth of fibrils.^68^ The creation of a nucleus encounters significant free energy barriers.^69–73^ Hence, successful nucleation events are extremely rare. The finding that the nucleation of p53 fibrils is hosted in the mesoscopic clusters suggests that the exaggerated p53 concentration in the clusters increases that probability of a fluctuation that overcomes the free energy barrier and evolves to a fibril nucleus. The proposed mechanism of fibrillization hosted and facilitated by mesoscopic clusters drastically deviates from the accepted sequential association of single solute molecules. Non-classical nucleation is a recently proposed mechanism of phase transformation that diverges from the canon of J.W Gibbs; it guides the assembly and defines the properties of numerous other protein solids such as crystals^22–25^ and sickle cell hemoglobin polymers.^74,75^

The clusters may represent a fibril-independent pathway to oncogenicity: unidentified pre-nuclear aggregates of two p53 mutants (R282W and R100P) were shown to accumulate the tumor suppressors p63 and p73.^14^ In this respect, the p53 liquid condensates may be akin to other protofibrillar assemblies known to trigger disease.^32^ Two-step nucleation of mutant p53 amyloids suggests means to control fibrillization and the associated pathologies through modifying the cluster behaviors. In addition, formation of clusters that combine mutant p53 with cancer suppressors (wild type p53, p63, p73, and others) may expedite the fibrillization of the suppressors into one-component fibrils or in fibrils that also incorporate mutant p53. The suggested two step mechanism of co-aggregation presents an alternative to the generally accepted templating pathway, which relies on a pattern provided by an existing fibril to guide the assembly of fibrils of a distinct protein. In a broader context, findings reported here exemplify interactions between distinct protein phases that activate complex physicochemical mechanisms operating in biological systems.

p53 R248Q exhibits enhanced cluster formation, faster fibrillization, and a richer variety of small oligomers compared to wild type p53. The demonstrated two-step nucleation of mutant p53 fibrils suggests that fibrillization may be faster owing to larger cluster populations whose extended volume enhances the probability of nucleation. The enhanced oligomerization is likely independently controlled. Wild type and p53 R248Q carry identical tetramerization domains. The distinct oligomer distributions of the two variants suggest that, in contrast to the native tetramers, the small oligomers are not due to interactions of the tetramerization domain, but likely to enhanced attraction between the DNA binding domains

The enhanced cluster formation of the R248Q mutant is not due to enhanced interaction in the disordered transactivation and proline-rich segments that are identical to those of wild type p53. Molecular models of the conformations of the wild type and p53 R248Q indicate that the enhanced cluster formation by the mutant is due to changes in the structured DNA binding domain that are promoted by the mutation and propagate allosterically to the N and C termini of the DBD. The shift of the N-terminus tail exposes the aggregation-prone motif ILTIITL (residues 251 to 257) and reinforces aggregation. The found role of the destabilization of the core domain in p53 condensation is consistent with the localization of most cancer-associated mutations in the structured DNA binding domain,^15^ and the role of mutant p53 aggregation in cancer.

## Online Content

Methods, along with additional Extended Data display items, are available in the online version of the paper; references unique to these sections appear only in the online paper.

## Data availability

The datasets generated during and/or analysed during the current study are available from the corresponding author on reasonable request.

## Acknowledgments

We thank Dominique Maes, Frederic Rousseau, and Joost Schymkowitz for valuable discussions of p53 aggregation, protein conformational stability, and protein expression and translocation, and the latter two for providing the cell lines used in this work. This work was supported by the National Science Foundation (Award Nos. DMR-1710354 and DMR-1705464), the National Institutes of Health (Award No. 1R21AI126215-01), CDMRP (Award No. CA160591), CPRIT (Award No. RP180466), MRA (Award No. 509800) and NASA (Award Nos. NNX14AD68G and NNX14AE79G). Additional support was provided by the Center for Theoretical Biological Physics at Rice University sponsored by NSF (Grant PHY-1427654)

## Author contributions

P.G.V., A.B.K., and N.V. conceived this work, P.G.V. and N.V. designed the experiments, D.Y. and M.S. expressed and purified wild type and mutant p53, D.Y. characterized p53 clusters and fibrils, A.S., M.F., and N.V. characterized aggregation in cell culture, A.D., A.K., and A.B.K carried out simulations of p53 structure. P.G.V., N.V., A.D., A.B.K. and M.C.B. wrote the paper, M.C.B. edited the text. All authors discussed the results and commented on the manuscript.

## Author information

The authors declare no competing financial interest. Readers are welcome to comment on the online version of the paper. Correspondence and request for material should be addressed to P.G.V.

## Methods

### Cell Culture

MCF7 (ATCC), a human breast adenocarcinoma cell line, and HCC70 (ATCC), a human breast carcinoma with mutant p53, were cultured in EMEM (Quality biological, 112-018-101) and RPMI (HyClone, SH3002701), respectively, supplemented with 10 % (v/v) fetal bovine serum (R&D systems, S11550), 1 % (v/v) HEPES (Corning, MT25060CI), 1 % (v/v) sodium pyruvate (Corning, 25-000-CI), 1 % (v/v) penicillin-streptomycin (Corning, 30-002-CI), and 1 % (v/v) L-Glutamine (Corning, 25-005-CI). Both cell lines were tested for mycoplasma and incubated in 37°C and 5 % CO_2_ incubator.

### Cell Treatment

20,000 HCC70 cells were seeded on one compartment of a 4-compartment CELLVIEW cell culture dish (Greiner, 627871). The next day, the cells were washed twice with PBS (HyClone, 16750-122) and 250 μl no-serum RPMI was added to the cells. The equal volume of 1,6-Hexanediol (Sigma, 240117, 6%) in no-serum RPMI was added on the petri dish and incubated for 30 seconds. Finally, the media was removed, and the cells were fixed immediately as described below.

### Immunostaining and ThT Staining

The cells were washed twice with PBS and fixed by incubating in 4% paraformaldehyde for 20 min at room temperature. Following three washes with PBS, the cells were then permeabilized and blocked by exposure to 0.5 % triton X-100 in PBS containing 2 % Bovine Serum Albumin for 30min at room temperature. The antibody staining was performed separately for Pab240 and DO-1 antibodies. For Pab240 staining, the cells were incubated with p53 antibody Alexa Fluor^®^ 546 (Santa Cruz, sc-99 AF546, 1:50 dilution, 4 μg/ml) in dark for one hour in room temperature. For DO-1 staining, the cells were incubated with p53 antibody Alexa Fluor^®^ 647 (Santa Cruz, sc-126 AF647, 1:50 dilution, 4 μg/ml) in dark overnight in 4°C. After washing 3 times with PBS, Thioflavin T (ThT, Sigma, T3516) staining, if necessary, was performed by adding 20 μM ThT in PBS in the dark for 15 min at 37°C. The cells were washed three times with PBS again and incubated with Hoechst 33342 (Sigma, 14533, 10 μg/ml) for 20 min in 37 C and washed twice with PBS before acquiring the images.

### Confocal microscopy

A Nikon (Minato, Tokyo, Japan) Eclipse Ti2 inverted microscope equipped with a 100x, Nikon, Plan Apo Lambda, oil, 1.45 NA objective was used for imaging. 3D images (z-stacks, 0.2 μm steps, ~60 slices) were taken from different field of views using DAPI, FITC, TXRed and Cy5 channels (Figures S1, S1, and S3).

### Analyzing Confocal Images

Z-stacks of 16-bit images were extracted for each channel and processed in ImageJ (National Institutes of Health (NIH), USA) using a series of plugins. First, images were cropped for single cells to assure the detected objectives belong to a single cell. Then, a series of methods including background subtraction, 3D watershed, 3D objective counter plugin,^1^ 3D ROI Manager plugin,^2^ and 3D Viewer were applied to construct the 3D images and measure the volume of p53 puncta (Figure S4).

### Bacterial Expression

Plasmid pET15b-TP 53, containing N-terminal 6-His-WT-p53 (1-303) (Addgene, 24859), was used to express p53. For p53-R248Q, a point mutation was introduced in pET15b-TP 53 by PCR and Gibson Assembly with two restriction enzymes (NdeI (New England Biolabs, R0111) and BamHI (New England Biolabs, R0136)) and four primers (forward primer 1: 5’-CACAGCAGCGG CCTGGTG-3’, forward primer 2: 5’-GCATGAACCGGAGGCCCATCCTCACCATCATCA-3’, reverse primer 1: 5’-TTCCTTTCGGGCTTTGTTAGCAGCCG-3’, reverse primer 2: 5’-AGTGTGATGATGGT GAGGATGGGCCTCCGGTTCAT-3’). The plasmids for p53-R248Q were transformed into Rosetta™ 2 (DE3) (Sigma Aldrich, 71397) by electroporation. After transformation, 50 μL of the solution was spread on a LB-agar plate with 100 μg/ml ampicillin (Fisher Scientific, BP1760) and incubated at 37 °C overnight. A single colony from the LB-agar plate with ampicillin was inoculated into 10 mL of media containing 20 g/L tryptone (Fisher Scientific, BP1421), 10 g/L yeast extract (Fisher Scientific, BP1422), 10 g/L NaCl (Fisher Scientific, S2711), and 100 μg/ml ampicillin. The culture was grown at 37 °C at 250 rpm. When optical density measured at 600 nm (OD_600_) reached 1.5 ~ 2.0, the culture was diluted as 1:100 with 1 L of same media with 0.2 mM ZnCl_2_ (Sigma Aldrich, 96468), and the culture was grown for 90 min at same condition such as culture with 10 ml volume. When OD_600_ was ~ 0.5, 1 L media with cells was placed in ice for cold shock. After 25 min, isopropyl β-D-1-thiogalactopyranoside (IPTG) (Denville Scientific Inc, CI8280) was added at 0.1 mM for induction. The culture was grown at 15 °C at 250 rpm overnight.^3–4^

### Cell Lysis

The cells were pelleted by centrifugation 5 °C at 4000 rpm for 1 hour by a Sorvall Legend X1R Centrifuge (Thermo Fisher) and resuspended in 20 mL of 100 mM KH_2_PO_4_/K_2_HPO_4_ (Fisher Scientific, AC263790010) with pH = 8.0, 300 mM NaCl, 5 % (v/v) glycerol (Sigma Aldrich, G7757), and 1 mM Tris(2-carboxyethyl)phosphine (Sigma Aldrich, C4706). 1 mL of 10X EDTA free protease inhibitor cocktail (SIMGA FAST protease inhibitor tablets) (Sigma Aldrich, S8830) was added to the cell suspension. The suspension was transferred into two 50 mL tubes to lyse the cells. Each aliquot was sonicated on ice four times for 30 seconds with 15 min intervals between each sonication with Q-SONICA MISONIX XL-2000 (11 watt output). After sonication, lysate was centrifuged at 5 °C with 13000 rpm for 15 min by Avanti J-E Ultra Centrifuge (Beckman Coulter). Supernatant from centrifugation was filtered with 0.45 μm SFCA syringe filters (Sigma Aldrich, CLS431220) prior to purification.

### Purification

10 mL of Ni sepharose™ Fast flow column (GE Healthcare Lifescience, 17531801) was equilibrated with 50 mM of binding buffer containing 100 mM KH_2_PO_4_/K_2_HPO_4_, 300 mM NaCl, 5 % glycerol, 25 mM imidazole (Sigma Aldrich, 792527), and 1 mM TCEP with pH 8.0. Filtered supernatant from the preceding centrifugation was loaded into column. p53-R248Q was eluted with linear gradient of imadazole. Two fractions with most p53-R248Q (based on SDS-PAGE) were collected. Fractions were diluted with 1:3 with binding buffer for heparin purification including 20mM KH_2_PO_4_/K_2_HPO_4_, 2 mM TCEP, and 5% glycerol at pH = 5.9 for HiTrap™ Heparin HP column (GE Healthcare Lifescience, 17040601). p53-R248Q is eluted with linear gradient of NaCl. Two fractions with most p53 R248Q were collected again for buffer exchange Figure S5a,b).

Before centrifugation to concentrate one of fraction, 1 M L-Arginine (Sigma Aldrich, A5131) was diluted into fraction as 1:10 ratio to prevent aggregation^5^. One of collections was centrifuged with 30000 MWCO Amicon Ultra Centrifugal Filtration Unit (Amicon, UFC803024). Concentrated fraction was mixed with another fraction and was loaded into PD-10 desalting column (GE Healthcare Lifescience, 17085101) for buffer exchange with incubation buffer containing 50 mM Tris, 150 mM NaCl, 5 mM TCEP, and 10% Glycerol at pH = 7.9. After buffer exchange, the R248Q solution was frozen in liquid nitrogen and stored in freezer at −80 °C. For experiments, protein solution was thawed in cold room with ice. Protein and incubation buffer were filtered with 0.22 μm Polyethersulfone (PES) (CELLTREAT Scientific Products, 229746) syringe filters. The solution concentration was determined by absorbance measurement using a Du 800 spectrophotometer (Beckman Coulter) and extinction coefficient *ε* = 0.763 mL * *mg*^−1^ * *cm*^−1^ at 280 nm^4,6^.

The identity of p53 was confirmed with western blotting. Primary antibody was p53 (DO-1) (Santa Cruz Biotechnology, sc-126) and secondary antibody was m-lgGκ BP-HRP (Santa Cruz Biotechnology, sc-516102). Amersham ECL Select Western Blotting Dection Reagent (GE Healthcare Lifescience, RPN2235) was used for illumination by Amersham imager (GE Healthcare Lifescience) to confirm binding of secondary antibody (Figure S5c).

### Oblique illumination microscopy

p53 R248Q was monitored with oblique illumination microscopy (OIM) known as Brownian microscopy or particle tracking.^7–10^ In this method, a green laser illuminates a thin solution layer at an oblique angle such that the incident beam avoids the lens of a microscope positioned above the sample (Figure 2a).^11^ This method enables the detection of nano- and microscale objects through light scattered at wavevectors of order μm^-1^. The scattered intensity is proportional to the sixth power of the scatterers’ sizes; thus, in a solution containing objects of varying size the scattering signal is dominated by larger particles. This feature makes this technique particularly well suited to characterize the size and number distribution of the aggregates that appear as bright cyan spots in OIM micrographs (Figure 2b). The spots are counted by a custom-made image package form Nano sight. The concentration of the observed aggregates is determined from the number of spots in a frame and the observed volume *V* = 120 × 80 × 5 *μm*^3^.^9,12^ OIM records the Brownian trajectory of each particles in the image plane (Figure S6a) and calculates diffusion coefficient from correlation between the mean squared displacement 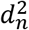 and the lag time Δ*t* (Figure 6b).^12^

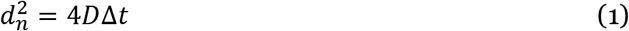

where *D* is the diffusion coefficient of the observed aggregate.

We used Nanosight LM10-HS microscope (Malvern Panalytical Inc) was used. Protein solution was loaded into a cuvette with volume about 0.3 mL and depth about 0.5 mm. The wavelength of illumination was 532 nm. Temperature was set between 15 and 42°C. Movies were acquired over 30 seconds. We found that objects recorded for times shorter than 1 s were interference spots from two or more clusters tracked for significantly longer times. This observation was supported by the estimate that a cluster with diffusivity D_2_ ≈ 10^-12^ m^2^s^-1^ would be detectable in a focal plane with depth 5 μm for about 25 s. Therefore, we excluded these short-duration objects from the determination of the cluster parameters. Five movies from distinct solution volumes were collected in each tested sample. The numbers of clusters in each range of sizes were averaged.

### Dynamic light scattering (DLS)

DLS data was collected by an ALV instrument (ALV-GmbH, Germany), which includes ALV goniometer, a He-Ne laser with wavelength as 632.8 nm), and an ALV-5000/EPP Multiple tau Digital Correlator. Normalized intensity correlation functions *g_2_*(*q*, *τ*) were collected at a fixed scattering angle of 90° for 60 seconds. The characteristic diffusion times *τ*_1_ and *τ*_2_ for monomers and aggregates were calculated by fitting the normalized correlation function with an exponential fit^13,14^:

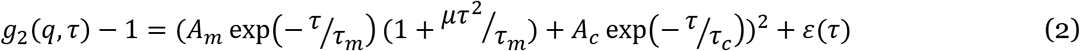

where *A*_1_ and *A*_2_ are amplitudes, which are proportional to the intensity scattered by the monomers and aggregates, respectively, and *ε*(*τ*) is background noise in the correlation function. From *τ*_1_ and *τ*_2_, the diffusivities of monomer and condensates, *D*_1_ and *D*_2_, are determined from the relations *D*_1_ = (*q*^2^ *τ*_1_)^−1^ and *D*_2_ = (*q*^2^ *τ*_2_)^−1^, where 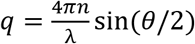 is the scattering wave vector at 90° with *λ* = 632.8 nm.

To calculate the average radii of monomers *R*_u_ and clusters *R*, we used the Strokes-Einstein relation, 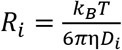 with *D*_1_ and *D*_2_. In this relation, *k_B_* is the Boltzmann constant, *T* is temperature, and η is the independently determined solution viscosity.^15^

### Static light scattering (SLS)

Osmotic compressibility was measured with the ALV instrument, discussed above. We measured the intensity scattered at 90° from p53 solutions with protein concentration varying from 0.12 – 0.4 mg mL^−1^ in incubation buffer: 50 mM Tris at pH = 7.85, 150 mM NaCl, 10% glycerol, and 5 mM TCEP. The average molecular weight *M_w_* and the second virial coefficient *B_2_* was calculated from the plot of *KC*_0_/*R_θ_* as a function of p53 concentration *C*_0_.^16,17^

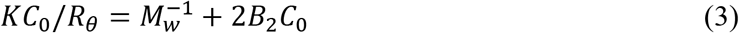

Here *R_θ_* = *I_θ_*/*I*_0_ is the Rayleigh ratio of scattered to incident intensity and *K* is the system constant, defined as 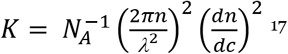, where *n* = 1.33 is the refractive index of the solvent and *dn*/*dc* = 0.20 is the first derivative of the refractive index with respect to protein concentration. *dn*/*dc* was measured using a Brookhaven Instruments differential refractometer operating at a wavelength of 620 nm.^18^

### ANS assays for characterization of p53-R248Q aggregation

The fibrillization kinetics of p53 R248Q was monitored at 37°C with 1-anilino-8-naphthalenesulfonate (ANS, Sigma Aldrich) assay^19–21^. The concentration of ANS was 18 mM in incubation buffer. The solution was filtered with 0.2 μm Teflon filter. The concentration of ANS was measured spectrophotometrically using extinction coefficient 18 mM^-1^ cm^-1^ at 270 nm^19^. Before loading sample, solution was kept on ice to prevent aggregation. Two exact identical samples were prepared and loaded in a 96-well plate. The fluorescence was excited at 350 nm, emitted at 500 nm, and recorded by an Infinite 200 PRO microplate reader (Tecan) at 10 minutes intervals over six hours.

Ficoll PM-70 (Sigma Aldrich) is used with the ANS assay. Stock concentration of Ficoll was 250 mg mL^-1^ with 200 mM NaCl in DI water for ionic strength of the buffer^15^. Concentration of p53-R248Q in incubation buffer with Ficoll was 6.5 μM to monitor growth of fibrils at 37 °C.

### Simulation and visualization for p53 Structures

The structure of wild type p53 was based on PDB structure 1TUP. We introduced the mutation R248Q in the 1TUP structure by choosing lowest score to optimize structure with mutation in Swiss-PdbViewer^22^. These images was made with VMD/NAMD/BioCoRE/JMV/other software support (http://www.ks.uiuc.edu/Research/vmd/). VMD/NAMD/BioCoRE/JMV/ is developed with NIH support by the Theoretical and Computational Biophysics group at the Beckman Institute, University of Illinois at Urbana-Champaign.^23^

### Coarse-grained simulations of the p53 DBD domain

Molecular dynamics (MD) simulations of DNA binding domain (DBD) of p53 were carried out using the AWSEM coarse-grained (CG) model.^24^ In this model, each amino acid is represented by 3 beads placed at the positions of C_α_, C_β_, and O atoms. The interactions between those beads are governed by a combination of physically motivated potentials, responsible for proper backbone geometry, secondary structure and long-range tertiary interactions, and bioinformatics terms that supplement the former potentials. Specifically, in this work, we employ the fragment memory potential that biases local (in sequence) conformations using experimentally solved structures containing similar motifs^24^ and evolutionary restraints inference from coevolutionary analysis of the target protein sequence^25^. The simulations were prepared using tools from the AWSEM-MD GitHub repository (https://github.com/adavtyan/awsemmd) based on the P53 DBD sequence (aa. 94-289). The evolutionary analysis was performed using RaptorX online server^26^, where contacts with a probability above 0.5 were incorporated in the model. The initial confirmation for MD simulations was based on PDB structure 1TUP, chain A, and simulations that were performed using the LAMMPS MD package^27^. The same setup was repeated for wild type and R248Q mutant of the DBD domain. In each case, the structure was minimized and equilibrated for 10 million time steps at 300K using the Nose-Hoover thermostat. The final state of the system (coordinates and velocities) was saved to perform any further simulations. In addition, simulations were performed for the truncated DBD domain (aa. 107-276, hereafter referred to as DBD core), where its N-terminal and C-terminal regions were removed.

### Free energy calculations

Starting from the equilibrated states of wild type and R248Q mutant of the DBD (as well as for the DBD core), we performed free energy calculations along the reaction coordinate *q*, which is defined as a similarity measure to a reference structure. Given the instantaneous coordinates *r* and the coordinates of the reference structurer *r*′, *q* can be defined as

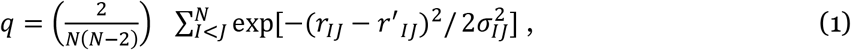

where *N* is the number of atoms in the structure and *σ_IJ_* is the standard deviation of the (*r_IJ_* − *r*′_*IJ*_ differences; *q* takes values from 0 to 1. The free energy calculations were performed using the Umbrella Sampling and the Weighted Histogram Analysis Method (WHAM)^28^. To perform the Umbrella Sampling simulations, a harmonic term *U* = *k*(*q* – *q*_0_)^2^ was added to the potential function. We used 243 windows, corresponding to different *k* and *q_0_* values (*k*=20,000, 50,000, and 100,000 kcal/mol, delta *q*=0.005), to calculate the free energy *F*(*q*) between *q*=0.3 and *q*=0.7. Further, 2D free energy profiles *F*(*q*, *Rg*) and *F*(*q*, *D*) along *q* and radius of gyration and *q* and end-to-end distance, respectively, were calculated using the reweighting technique. Each Umbrella Sampling simulation was run for 50 million time steps. Chain A of PDB 1TUP was used as the reference structure.

### Analysis and structural clustering

The structural diversity of p53 DBD was studied by clustering conformations near the local minimums of the 2D free energy profiles *F*(*q*, *R_g_*) and *F*(*q*, *D*). The clustering was performed using the Hierarchical clustering algorithm^29^. Consequently, several distinct conformations of DBD were identified for both wild type and the R248Q mutant (see Results section).

## Supplementary Figures

**Figure S1.**
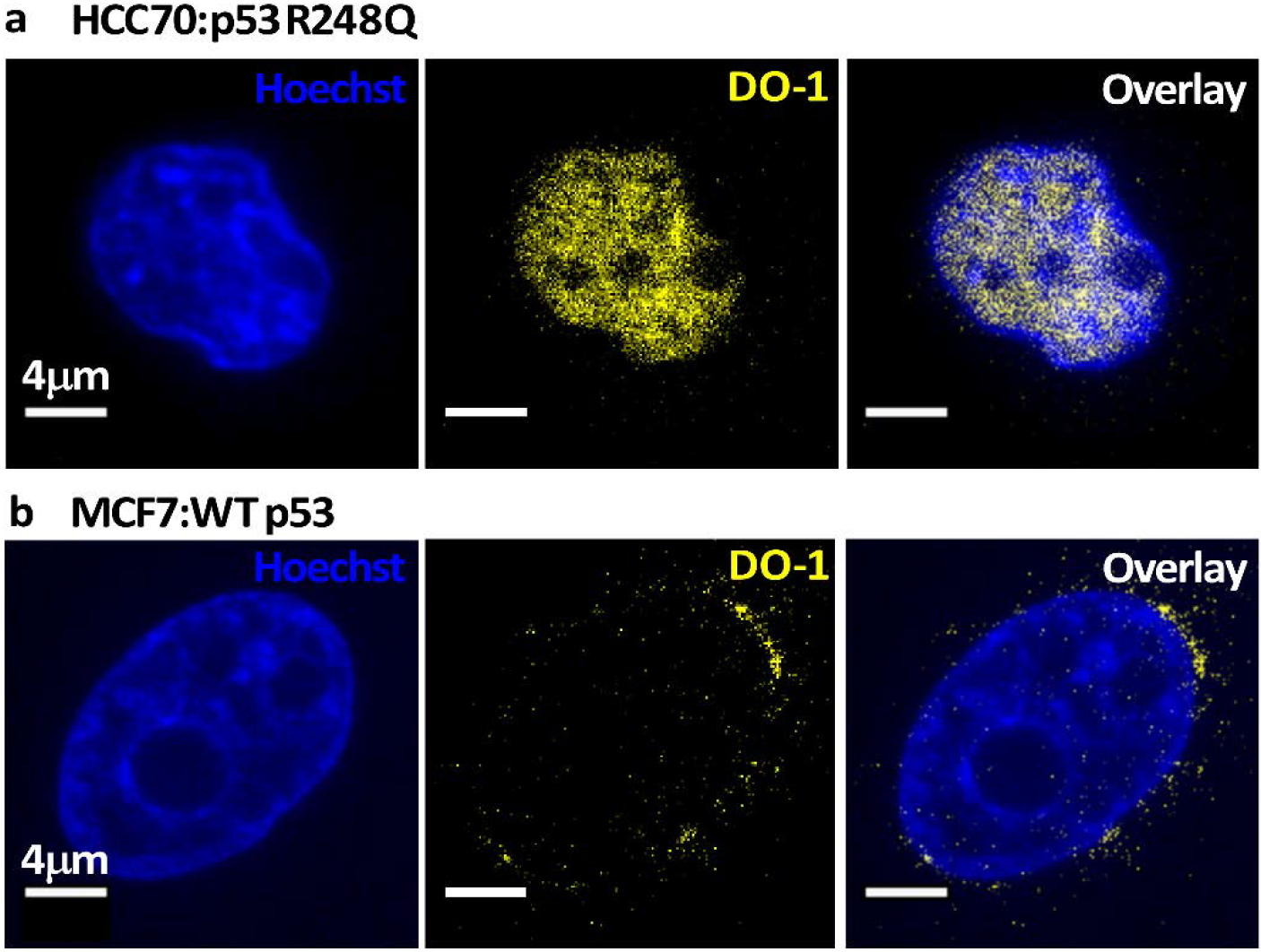
Unaggregated wild type p53 and p53 R248Q in breast cancer cells. **a,b.** Combined staining with Hoechst, which stains the nucleus, and DO1, which binds to unaggregated p53. Confocal images of **a.** A representative HCC70 single-cell. **b.** A repsreeitnative MCF7 cell.

**Figure S2.**
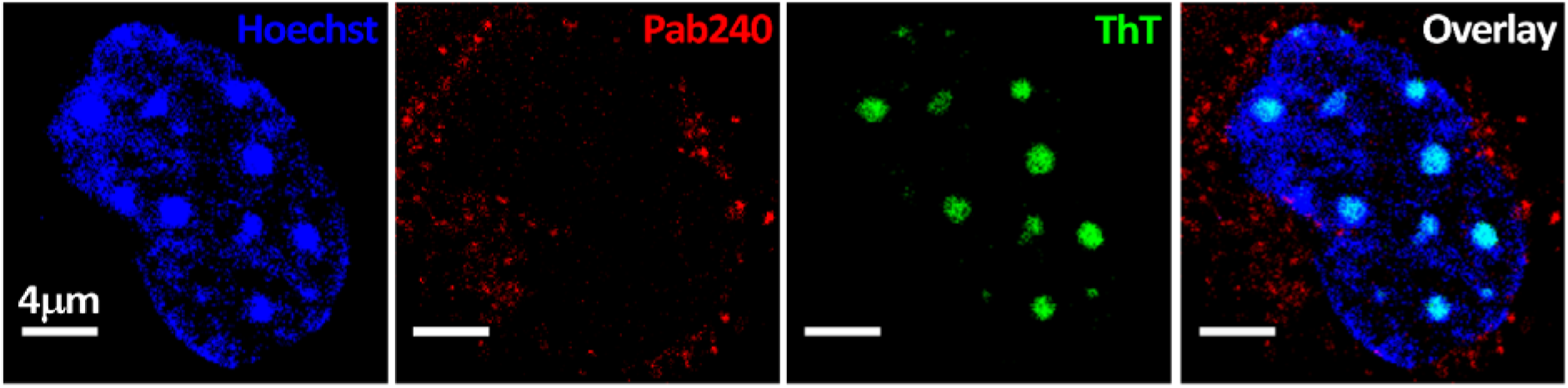
Aggregated p53 R248Q in a breast cancer HCC70 cell. Combined staining with Pab240, which binds to unfolded or aggregated p53, and ThT, which detects amyloid structures. The cell nucleus is stained with a Hoechst dye. A confocal image of a representative single cell is shown.

**Figure S3.**
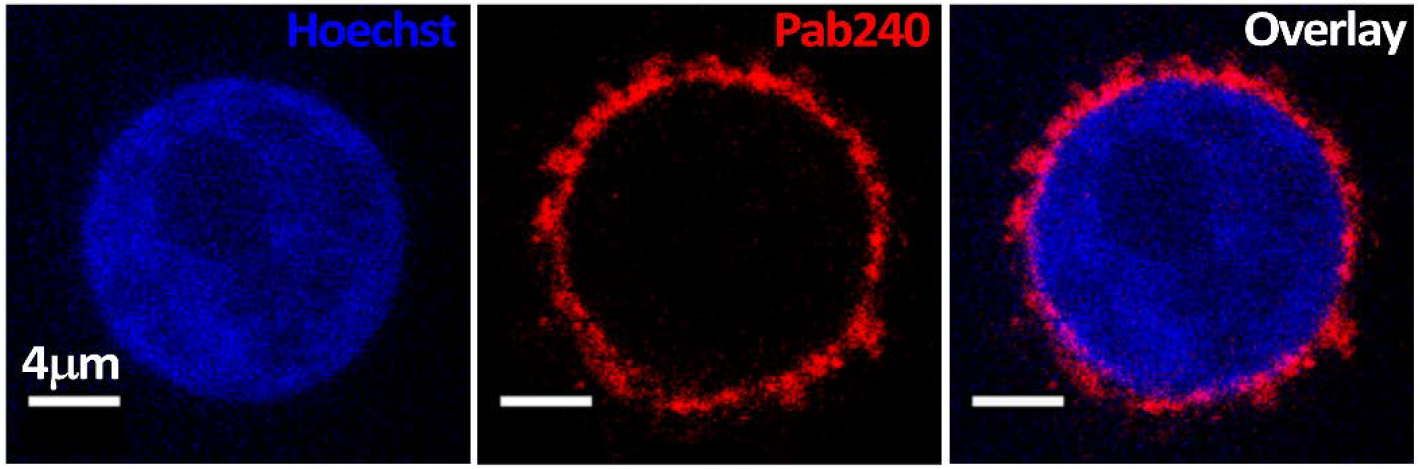
Aggregated/misfolded wild type p53 in a breast cancer MCF7 cell. Combined staining with Pab240, which binds to unfolded or aggregated p53, and Hoechst, which stains the nucleus. The confocal image of a representative single cell is shown.

**Figure S4.**
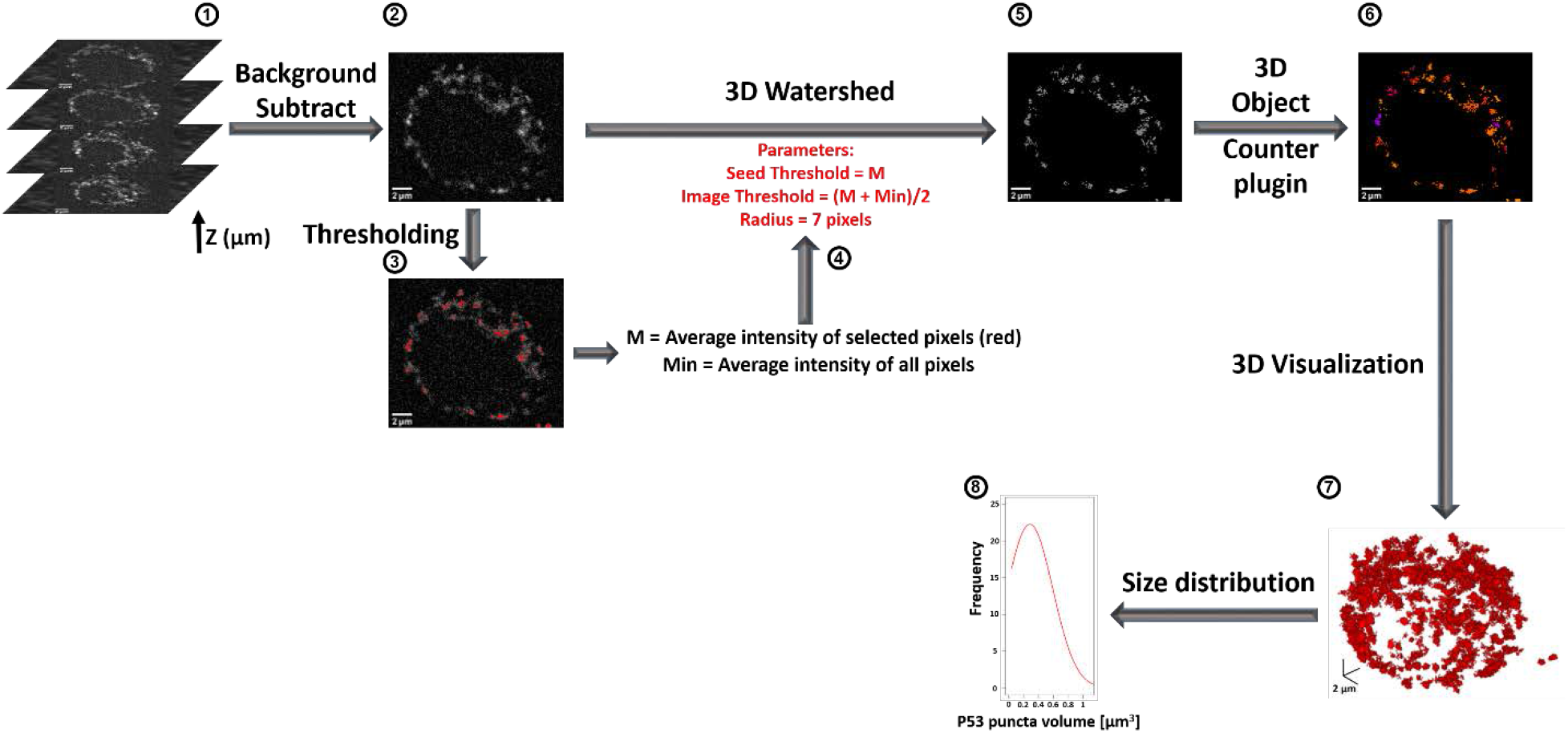
The workflow of image analysis for measurement and 3D construction of p53 puncta in a single cell. 1. Extract and crop Z stack images. 2. Subtract background across the stacked image. 3. Select p53 puncta by using intermodes thresholding to select pixels with high fluorescence intensity. 4. Calculate M as the average intensity of selected pixels and Min as average intensity across image. 5. Apply 3D watershed on the background-subtracted image using parameters calculated by M and Min. 6. Construct the 3D objects using the 3D object counter. 7. Construct the 3D image directly from objects detected from 3D object counter using the 3D Viewer plugin. 8. Plot the size distribution of p53 puncta for every single cell.

**Figure S5.**
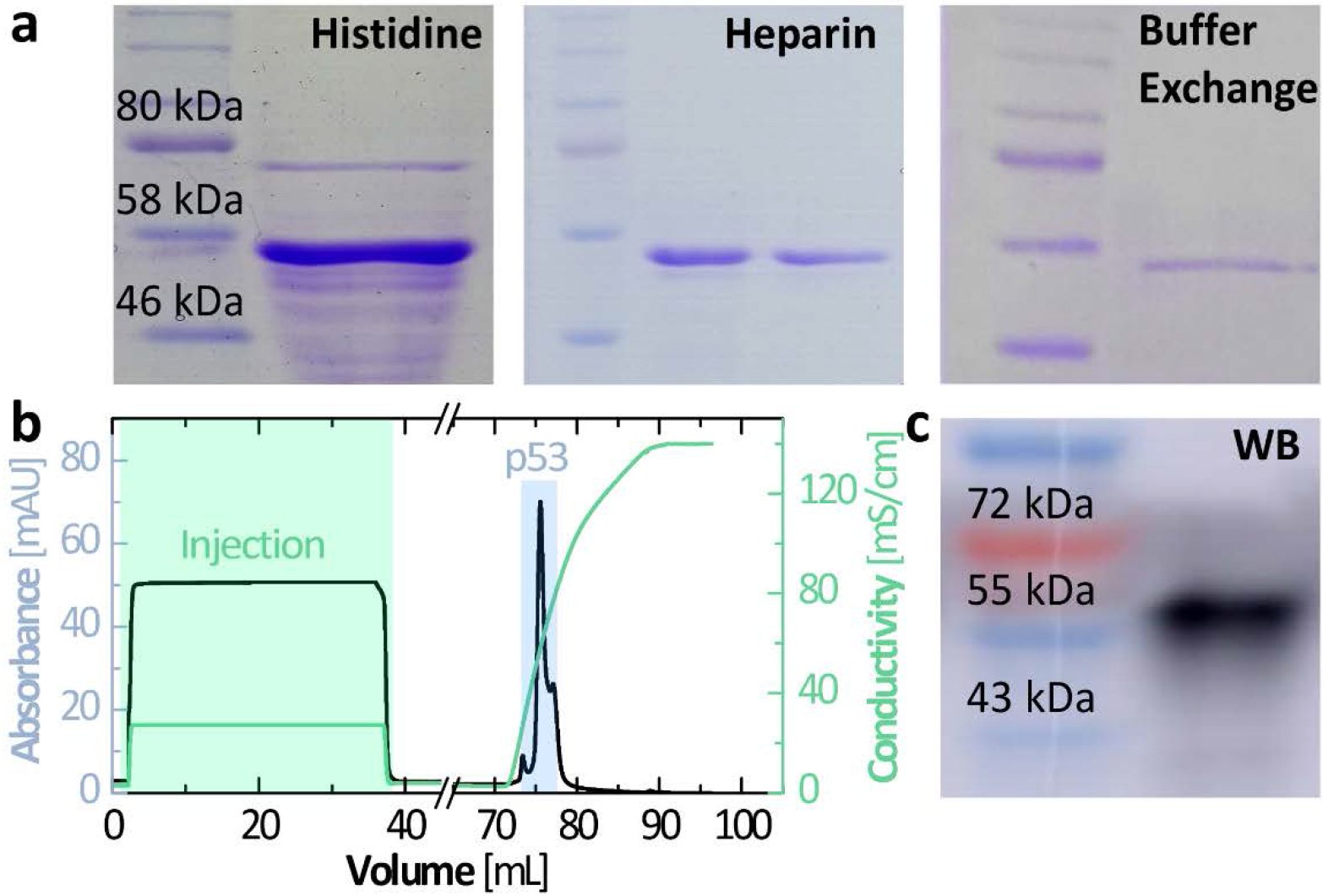
Purification and identification of p53 R248Q. **a.** SDS-PAGE after histidine purification (left), Heparin purification (middle), and buffer exchange (right). **b.** Absorbance at 280 nm and conductivity of a p53 R248Q solution from heparin resin chromatography column with 2 M NaCl gradient. **c.** Western blot of p53 R248Q (Primary antibody: DO-1; secondary antibody: m-lgGκ BP-HRP).

**Figure S6.**
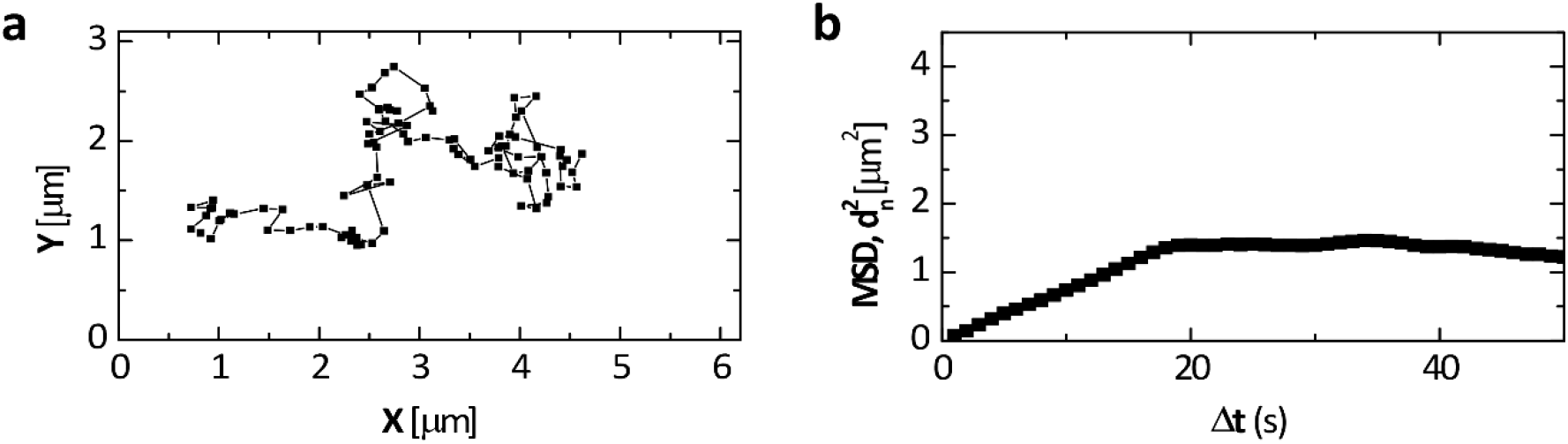
Oblique illumination microscopy. **a.** A trajectory of one particle determined by the OIM software from a sequence of images. **b.** Mean squared displacement *d^2^* as function of the corresponding lag time Δ*t*.

